# MuCor: Mutation Aggregation and Correlation

**DOI:** 10.1101/022780

**Authors:** Karl W. Kroll, Ann-Katherin Eisfeld, Gerard Lozanski, Clara D. Bloomfield, John C. Byrd, James S. Blachly

## Abstract

**Motivation:** There are many tools for variant calling and effect prediction, but little to tie together large sample groups. Aggregating, sorting, and summarizing variants and effects across a cohort is often done with *ad hoc* scripts that must be re-written for every new project. In response, we have written MuCor, a tool to gather variants from a variety of input formats (including multiple files per sample), perform database lookups and frequency calculations, and write many report types. In addition to use in large studies with numerous samples, MuCor can also be employed to directly compare variant calls from the same sample across two or more platforms, parameters, or pipelines. A companion utility, DepthGauge, measures coverage at regions of interest to increase confidence in calls.

**Availability:** Source code is freely available at https://github.com/blachlylab

**Supplementary data:** Supplementary data, including detailed documentation, are available online.

## 1 Introduction

Examination of genomic variants from multiple samples is a common procedure in bioinformatics. Whether to use existing tools or custom scripts depends on many factors, including the number of samples, source(s) of data, and scope of the project. A typical workflow may involve variant annotation, extraction of variants from VCF files, binning loci into features, calculating summary statistics, filtering, limiting by region, and writing to a tab-separated values table. Finally, this table is imported into a statistical package for calculations or to Microsoft Excel for use by end-users or inclusion as a data supplement to a paper.

Existing tools provide only parts of this workflow. The Genome Analysis Toolkit (McKenna et al., HYPERLINK \l “bookmark2” 2010) function `CombineVariants` and vcftools’ `vcf-merge` can merge records from VCF files, but produce only another VCF file in turn. The resultant VCF is necessarily focused on individual genomic locations, and a study of features (e.g., genes) requires additional work on the part of the bioinformatician. Further, a multi-sample VCF file may be viewed interactively in a genome browser but is not suitable as a locus-wise frequency table for the biologist or statistician. Transformation of such a VCF to a detailed report again requires additional processing. Some commercial products purport to be end-to-end solutions, but these are not available to all.

Motivated by these factors, we developed MuCor, a flexible tool for variant aggregation and summarization. MuCor collects variants from an arbitrary number of sources, assigns them to user-defined features (typically but not necessarily genes), looks them up in variant databases, calculates metrics, and generates reports, all within a configurable and extensible framework.

## 2 Implementation and Usage

### 2.1 Implementation

MuCor is written in Python (≥2.7.0) and uses the following libraries not included with the Python standard library: `numpy` (van Der Walt et al., HYPERLINK \l “bookmark3” 2011), `pandas`, and `HTSeq` (Anders et al., HYPERLINK \l “bookmark1” 2015). It can optionally make use of `pytabix` and `xlsxwriter` to enable additional functionality. MuCor is used in two stages: *setup* and *run*.

Project setup can be as simple as running the configure script, yielding a JSON settings file. In ‘autodetect’ mode, the script scans the project directory recursively for all supported variant call files belonging to a supplied list of sample IDs; a single sample may have more than one associated VCF file (e.g., if SNV and indel detection are performed separately). The setup script also establishes links to variant databases against which the study samples will be checked. The final configuration can be adjusted prior to run phase.

In the run phase, MuCor parses the JSON configuration file and reads samples, databases, annotation, and region files in to the analysis core. It then groups variants according to features specified in the annotation, optionally limited to region(s) of interest, calculates summary metrics, and reports the information about each variant, feature, and sample in output reports. The key vehicle by which this is accomplished is the pandas DataFrame (df). By leveraging the grouping, aggregation, and pivoting functions native to pandas, the variant data can be swiftly manipulated in myriad ways. For example, all variants can be condensed to the gene level using `df.groupby`, while data could be reformatted into a sample × gene matrix with `df.pivot` or `df.stack`.

The three steps of the run phase—input, aggregation and analysis, and reporting—are distinctly separated to make MuCor easy to extend. New input and output formats are easily coded (see supplementary text), while any run can be limited to regions of interest and cross-referenced against an arbitrary number of variant databases with merely a configuration change. Figure 1 summarizes MuCor’s modular approach.

**Figure 1:**
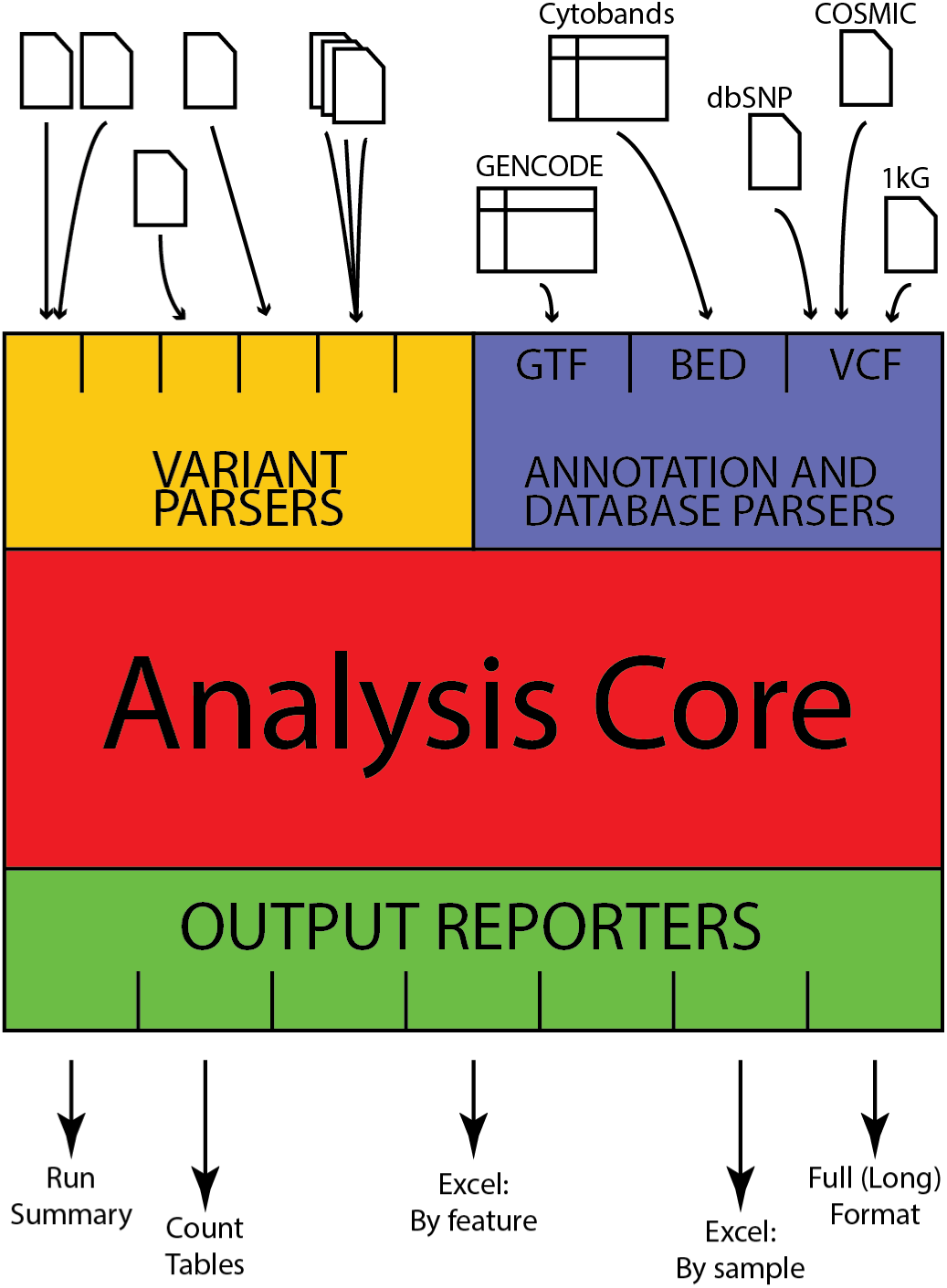
Schematic of MuCor’s inputs, engine, and outputs. Many more input and output types are available. See supplementary materials.

MuCor is not a variant annotator: functional effect prediction is better left to specialized tools. In support of this, MuCor accepts as input data that have already been decorated with a functional effect prediction, retains this information through processing, and passes it through to the output. Furthermore, some MuCor reports are in a form suitable for further processing by variant effect prediction software, a more efficient approach for large cohorts (i.e., hundreds to thousands of samples) than per-sample effect prediction.

### 2.2 Usage

Detailed instructions and sample workflows are given in the supplementary material. Configuration begins by specifying an annotation in GTF/GFF3 format, a feature type for grouping, a list of samples, and one or more report types. The user may optionally specify reference databases (e.g., dbSNP, 1000 Genomes, COSMIC) and limit analysis to regions of interest. Variant files are automatically detected. The resultant JSON file may be hand-edited prior to execution, or retained for reproducibility.

A corresponding run requires only passing the configuration file; output is written to a prespecified directory. Reports range in scope from summary-level counts across broad regions of interest to variant-specific metrics on a per-sample basis. The supplementary material contains a list of all report types and some example reports.

Finally, DepthGauge, a companion program, queries source BAM files for total reads at each defined region in all samples to improve confidence in the veracity of wild-type calls.

The generality of MuCor makes it useful for additional applications beyond comparing variants across samples. We have also used it to compare sequencing platforms, variant calling tools, and pipelines within a single sample. In this case, the comparisons reveal concordance and discordance between platforms or tools, rather than between samples as in the canonical usage. See example workflow 3 in the supplementary text.

## 3 Conclusions

We have created MuCor, a tool to aggregate and report genomic variants in configurable, meaningful groups. A key goal is generality, that it may replace the *ad hoc* scripting often performed in the course of routine bioinformatic analyses of multiple samples. A flexible runtime and modular architecture ensure broad applicability: we have applied MuCor for its intended use by aggregating hundreds to thousands of cases, but also for novel uses, such as comparing different variant calling software pipelines or different sequencing platforms. By separating input parsers, the annotation and analysis core, and output reporting into distinct components, MuCor is modular and expandable through new input plugins, new annotations or auxiliary databases, and definition of new output report formats.

## Acknowledgement

### Funding

This work was supported by the National Institutes of Health [P30 CA016058] and in part by an allocation of computing resources from The Ohio Supercomputer Center.

### Conflict of interest

None declared.

